# The relative impact of evolving pleiotropy and mutational correlation on trait divergence

**DOI:** 10.1101/702407

**Authors:** Jobran Chebib, Frédéric Guillaume

## Abstract

Both pleiotropic connectivity and mutational correlations can restrict the decoupling of traits under divergent selection, but it is unknown which is more important in trait evolution. In order to address this question, we create a model that permits within-population variation in both pleiotropic connectivity and mutational correlation, and compare their relative importance to trait evolution. Specifically, we developed an individual-based, stochastic model where mutations can affect whether a locus affects a trait and the extent of mutational correlations in a population. We find that traits can decouple whether there is evolution in pleiotropic connectivity or mutational correlation but when both can evolve then evolution in pleiotropic connectivity is more likely to allow for decoupling to occur. The most common genotype found in this case is characterized by having one locus that maintains connectivity to all traits and another that loses connectivity to the traits under stabilizing selection (subfunctionalization). This genotype is favoured because it allows the subfunctionalized locus to accumulate greater effect size alleles, contributing to increasingly divergent trait values in the traits under divergent selection without changing the trait values of the other traits (genetic modularization). These results provide evidence that partial subfunctionalization of pleiotropic loci may be a common mechanism of trait decoupling under regimes of corridor selection.

## Introduction

One of the central problems in evolutionary biology is understanding the processes through which new traits arise. One process that can lead to the creation of new traits is when existing traits become differentiated from one another because they are selected for a new purpose (Rueffler et al., 2012). There has long been evidence that this can happen through gene duplication followed by trait decoupling (Muller, 1936; Ohno, 1970; Rastogi and Liberles, 2005; Han et al., 2009). One example in vertebrates is the differentiation of forelimbs from hind limbs, where the same gene that was responsible for both fore and hind limb identity in development diverged (Graham and McGonnell, 1999; Minguillon et al., 2009; Petit et al., 2017). In this case, the paralogous genes *Tbx4/Tbx5* that encode transcription factors for fore/hindlimb identity likely evolved from the same ancestral gene, and their expression differentiated after duplication (Minguillon et al., 2009). Somehow during selection for functional divergence, there was a decoupling of genetically integrated traits, which allowed them to respond to selection as independent genetic modules (Wagner and Altenberg, 1996; Hansen, 2006). Genetic decoupling was likely also responsible for the evolution of trait divergence in vertebrate metameric segmentation into differentiated somites, and the emergence of cell differentiation in multicellular organisms (Holley 2007; Wagner et al. 2019; but see Newman 2020).

Although modular structures in phenotypic covariation (where phenotypic variation is more correlated within groups of traits than between them) are found in a wide range of organisms, including yeast, round worms, mice, and humans (Jiang et al., 2008; Wang et al., 2010; Hlusko, 2016), the underlying genetic architectures producing genetic integration between traits are still uncertain. Genetic integration, constraining the decoupling of traits, may arise from pleiotropic connections between loci and traits, where they may or may not create genetic and phenotypic covariation (Baatz and Wagner, 1997; Kenney-Hunt et al., 2008; Smith, 2016). When selection favours the divergence of traits, the constraining effect of pleiotropy may come in two forms: a pleiotropic connectivity effect **or** a mutational correlation effect (Stern, 2000). A pleiotropic connectivity effect depends on how highly pleiotropically connected a gene is. For instance, a gene product (e.g. enzyme, transcription factor, etc.) may affect more than one trait (or function) by having multiple substrates or binding sites, thus affecting multiple downstream processes. This may constrain the evolutionary divergence of those traits because the effect of a mutation beneficial for one trait may be deleterious for other traits (when those other traits are under stabilizing selection). It is expected that the net fitness effect of a pleiotropic mutation is decreased in proportion to the number of traits it affects (Orr, 2000; Welch and Waxman, 2003; Martin and Lenormand, 2006). Therefore, a pleiotropic connectivity effect can constrain divergent trait evolution even without creating genetic correlation among traits (a.k.a, hidden pleiotropy Wagner, 1989; Baatz and Wagner, 1997). Whereas, a mutational correlation effect is the effect of a mutation affecting how correlated are the effects of mutations at pleiotropic loci. Thus, a mutational correlation effect may induce correlated changes in the traits affected by pleiotropic loci. However, the strength of the correlational effect of the mutations is not dependent on the number of traits affected but on the properties of the genes, processes, or traits affected. When those effects are correlated among traits, they can constrain trait decoupling in addition to those caused by the dimensionality of the pleiotropic loci (Lande, 1979; Arnold, 1992; Stern, 2000).

Biological examples may help to illuminate the distinction between the two types of pleiotropy that can hinder the decoupling of traits. Imagine a transcription factor (TF) that has multiple target binding sites affecting the expression of multiple genes, which in turn affect several traits. If the binding sites have identical sequences, then mutations in the gene encoding the TF are expected to be in perfect correlation with respect to their effects on the traits. In this scenario, a mutational correlation mutation may be a mutation in one of the binding sequences that leads to differential binding of the transcription factor (Figure 1). Now, that the binding sites are no longer identical, mutations in the gene encoding the TF may no longer have perfectly correlated effects on the traits. As the name suggests, the mutational correlation mutation has affected the correlation between effects of mutations in the TF’s gene on the traits it affects. Whereas, another type of mutation might affect a TF’s access to one of it’s binding sites (e.g. by methylating the DNA in the region of that binding site). If this type of mutation causes the TF to affect more or less traits than it did before the mutation, it would be considered a pleiotropic connectivity mutation.

**Figure 1:**
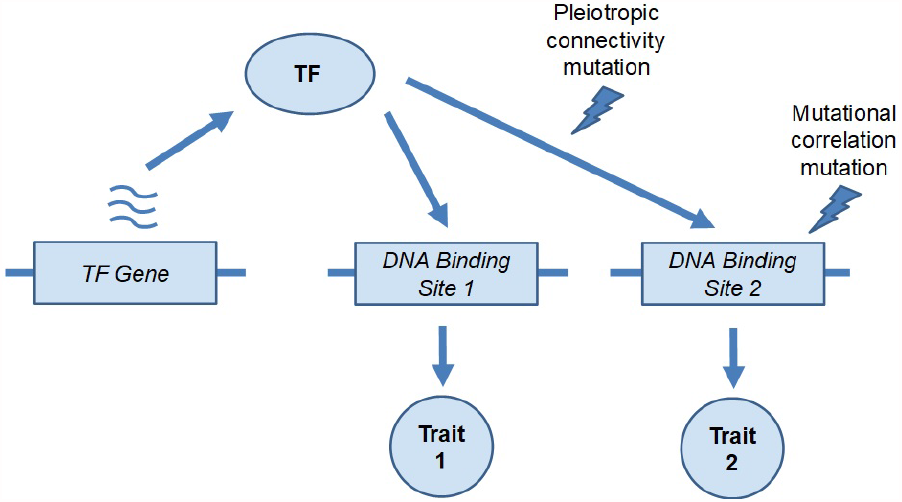
The two types of mutations affecting pleiotropic effects using a transcription factor (TF) as an example.

Both pleiotropic connectivity and mutational correlations can evolve as a result of divergent selection and affect the ability of traits to decouple from one another. Although previous models have included either evolution in pleiotropic connections or mutational correlation, their relative importance in constraining trait decoupling remains to be seen (Jones et al., 2003, 2007; Melo and Marroig, 2015; Chebib and Guillaume, 2017). Here we attempt to answer this question by using stochastic simulations, where individuals in a population can vary in both pleiotropic connections and mutational correlations, while applying divergent selection on some traits but not others, and then observing what affects the decoupling of traits.

## Methods

### Simulation model

We modified the individual-based, forward-in-time, population genetics simulation software Nemo (v2.3.46) (Guillaume and Rougemont, 2006) to allow for the evolution of pleiotropic connectivity and mutational correlations. The simulations consisted of a single population of size *N* of randomly mating, hermaphroditic, diploid individuals, with a probability 1*/N* of self-fertilization, similar to a classical Wright-Fisher population model. Each individual had two pleiotropic QTLs affecting four traits. The phenotypic value of each trait, *z*_*i*_ , was calculated by adding the allelic values at the two 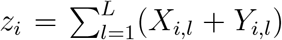 , where *X* is the maternally inherited allele and *Y* the paternally inherited allele, *i* is the trait number (*i ∈* [1, 4]), and *L* is the locus number (*L ∈* [1, 2])) (Figure 2). For simplicity, we assumed no environmental variance (i.e. heritability is 1).

**Figure 2:**
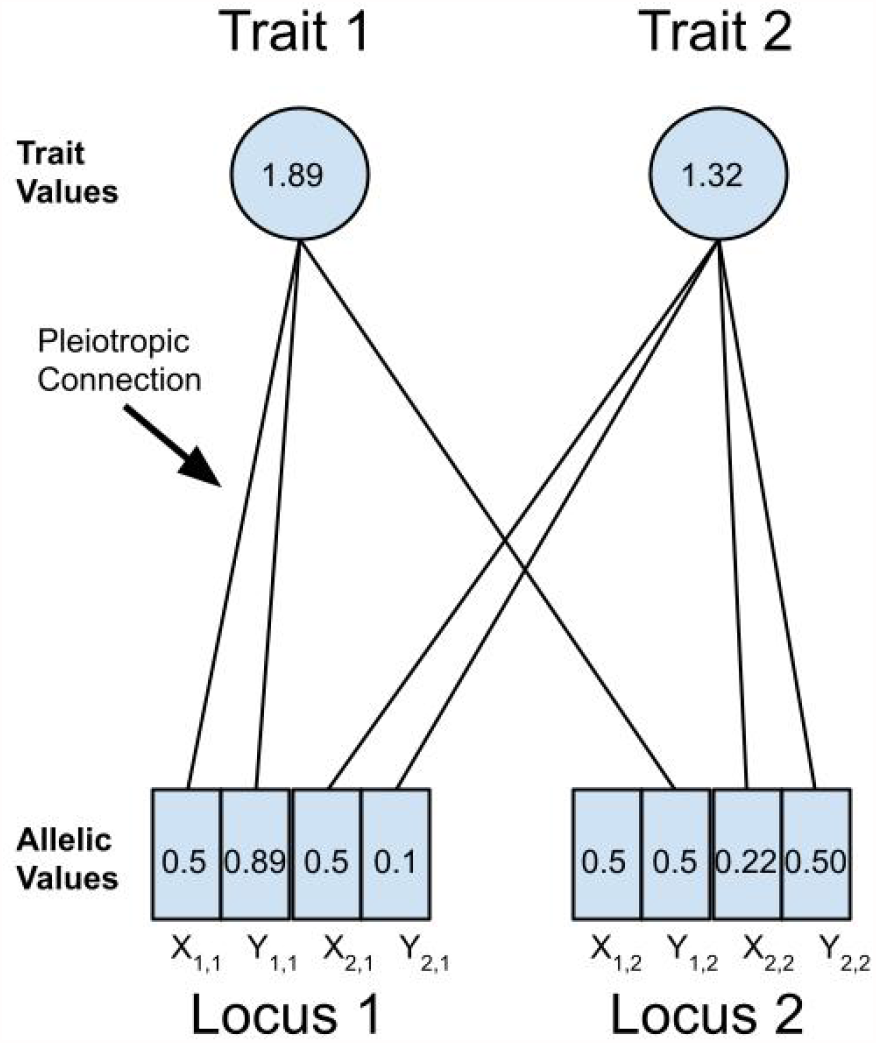
Pictorial representation of quantitative trait locus (QTL) alleles and the traits they affect with example values for illustration. Here, Trait 1 has a value of 1.89 determined by the sum of allelic values (*X*_1,1_, *Y*_1,1_, *andY*_1,2_) pleiotropically connected to it from Locus 1 (0.5 + 0.89) and Locus 2 (0.5), where *Y*_1,2_ represents the maternally inherited allele of Locus 2 that affects Trait 1. Trait 2 is affected differently by the two loci and has a value of 1.32 (0.5 + 0.1 + 0.22 + 0.5). The allelic values of a QTL were affected by mutation at a rate of *µ*. The pleiotropic connections between a QTL and a trait could be removed or added by mutation at a rate of µ_*pleio*_, and toggled whether an allelic value was added to a trait value or not.

Generations were non-overlapping and consisted of three main stages: mating, viability selection, ageing. In the mating stage, pairs of individuals were chosen to produce offspring (with a mean fecundity of three offspring to ensure population size replenishment). It was during the mating stage that recombination between loci and mutations occurred. In the viability selection stage, Gaussian stabilizing selection was applied on offspring and determined the survival probability of individuals, whose fitness was calculated as 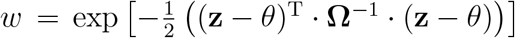 , where **z** is the individual trait value vector, *θ* is the vector of local optimal trait values, and **Ω** is the selection variance-covariance matrix (*n* × *n*, for *n* traits) describing the multivariate Gaussian selection surface. The **Ω** matrix is a diagonal matrix with diagonal elements corresponding to the strength of selection, *ω*^2^, on each trait (where strength of selection scales inversely with *ω*^2^), and off-diagonal elements corresponding to the strength of correlational selection, *ρ*_*ωij*_ , between traits *i* and *j*. In the ageing stage, the adults were removed from the population and the offspring matured into breeding adults for the next generation.

Three types of mutations (each with a separate mutation rate) were possible: mutations at the additive QTL affecting the traits, mutations at a set of modifier loci separately affecting the correlation of the mutational effects at the additive QTL, and mutations affecting the connectivity of the QTL to the traits, reducing or increasing the pleiotropic degree (the number of traits a locus affects) of each locus. The first type of mutations changed the allelic values of a QTL by randomly drawing effects from a multivariate normal distribution, with variance-covariance matrix **M**. The mutational effects were added to the existing allelic values at the QTL (continuum-of-alleles model; Crow and Kimura, 1964). These mutations appeared at a rate given by *µ*. The variance of the mutational effect for all traits were constant and set at *a*^2^ = 0.1 in the diagonal of the **M**-matrix. Each pairwise trait covariance of the **M** matrix was governed by its own separate modifier locus. We will refer to variance-standardized covariance values, or mutational correlation *ρ*_*µij*_ as the off-diagonal elements of the **M** matrix. As **M** is a 4 × 4 symmetrical matrix, the 6 *ρ*_*µij*_ coefficients were controlled by 6 diploid modifier loci, carried by each individual and inherited in the same manner as the additive QTL. Each individual thus carried its own **M** matrix. The second type of mutation thus changed these mutational correlation allelic values by randomly drawing from a uniform distribution (*-*0.2 * *log*[1 *-* **U**(0, 1)]), and adding the effect to the existing allelic value (which was bound between -1 and 1). These mutations appeared at a rate given by *µ*_*mutcor*_ . In order to get a particular mutational effect correlation, *ρ*_*µij*_ , the two mutational correlation allelic values of the corresponding modifier locus were averaged together (Figure 3). All the pairwise mutational effect correlations (*ρ*_*µij*_) were combined with mutational effect variances 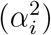 to create the **M** matrix for an individual, whenever a mutational effect on a QTL that directly affected traits was required. The third type of mutation affected the pleiotropic connections between QTLs and traits, determining whether the allelic value of a QTL was added to a trait value. A mutation of this type affected the pleiotropic connections between a trait and the maternally or paternally inherited alleles separately. Thus, QTLs could be ‘heterozygotes’ in their pleiotropic degree depending on the pleiotropic degree of the maternally and paternally inherited alleles. These mutations appeared at a rate given by *µ*_*pleio*_ (Figure 2).

**Figure 3:**
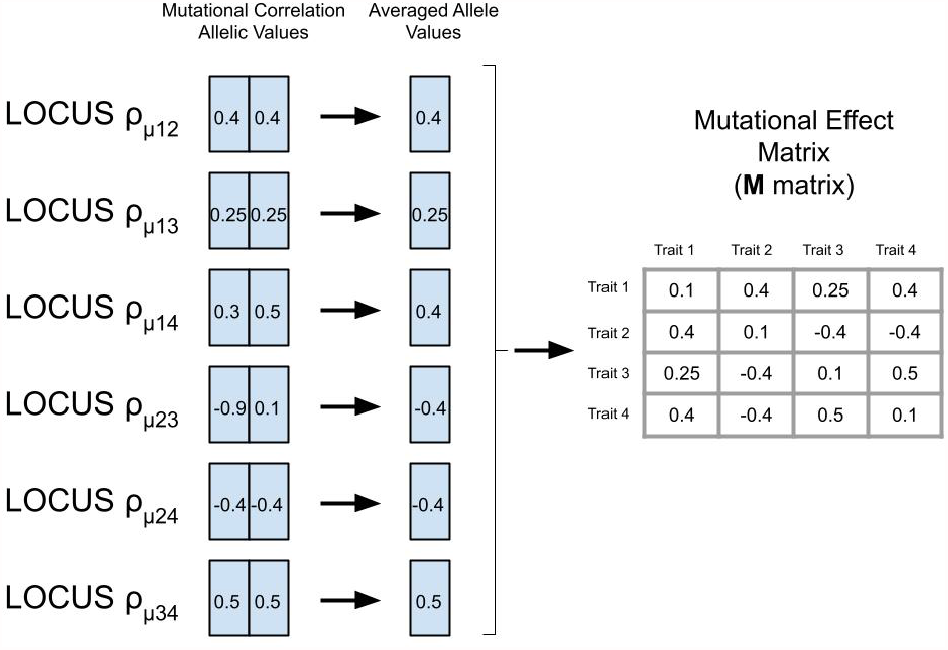
Pictorial representation of the modifier loci that contained the allelic values for producing the pairwise mutational effect correlations (*ρ*_*µij*_) between Traits *i* and *j*. The allelic values of a modifier locus were affected by mutation at a rate of *µ*_*mutcor*_ , and were averaged together to produce the corresponding correlation for the ***M*** matrix. The mutational effect variances, *α*^2^, remained static with a value of 0.1 for all traits.

### Experimental design

To understand the impact of divergent selection on the structure of genetic architecture, simulations were run with a population of 500 individuals that had two additive loci underlying four traits (Figure 4). The initial conditions were set to full pleiotropy (each locus affecting every trait) and strong mutational correlations between trait pairs (*ρ*_*µ*_ = 0.99). This way, mutational effects in phenotypic space were highly constrained to fall along a single direction, and reducing variation for divergent selection. All traits had an initial phenotypic value of 2 with equal allelic values of 0.5 at each allele of the two QTL.

**Figure 4:**
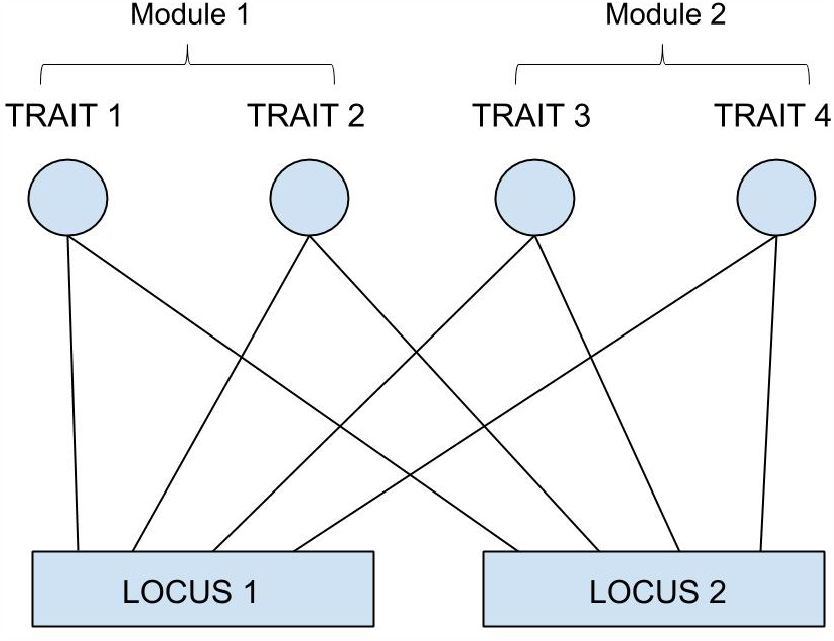
Pictorial representation of the genetic architecture modelled within individuals at the start of the simulations, with 2 loci, 4 traits, and full pleiotropic connectivity between them.

Selection regimes were designed to mimic divergent selection between trait modules, where Trait Module 1 included Traits 1 and 2, and Trait Module 2 included Traits 3 and 4. Initially, optimum trait values, *θ*_*k*_ , (*k ∈* 1, 2, 3, 4), were all set to 2 (the same as the initial trait values). There was moderately strong stabilizing selection on each trait (*ω*^2^ = 5), strong correlational selection between traits in the same trait module (*ρ*_*ω*12_ = *ρ*_*ω*34_ = 0.9), and no correlation between traits in different trait modules (*ρ*_*ω*13_ = *ρ*_*ω*14_ = *ρ*_*ω*23_ = *ρ*_*ω*24_ = 0). After this, divergent directional selection proceeded by maintaining constant optimal trait values for Traits 3 and 4 (*θ*_3_ = *θ*_4_ = 2) and increasing the optimal trait values for Traits 1 and 2 by 0.001 per generation for 5000 generations, bringing the trait optima to *θ*_1_ = *θ*_2_ = 7 (corridor model of selection *sensu* Wagner, 1984; Bürger, 1986). These 5000 generations of divergent, directional selection on Traits 1 and 2 were then followed by 5000 generations of purely stabilizing selection.

In order to compare the differential effects of evolving pleiotropic connectivity and evolving mutational correlations on trait decoupling, nine different simulations were run with all combinations of three different rates of mutation in pleiotropic connectivity and mutational correlations (*µ*_*pleio*_ and *µ*_*mutcor*_ = 0, 0.001, or 0.01) representing no evolution, and mutation rates below, at and above the QTL allelic mutation rate (*µ* = 0.001), respectively.

Simulations were also run with initial mutational correlations between all pairs set to 0 (*ρ*_*µ*_ = 0) to compare highly constraining genetic architecture (within a corridor selection regime) to ones with no constraints in the direction of mutational effects.

We observed general patterns of average trait value divergence, population fitness, genetic correlation, pleiotropic degree (the number of traits a locus affects) and mutational correlation. In the case of pleiotropic degree, the two loci affecting trait values were sorted into a high and low pleiotropic degree locus for each individual before averaging over populations or replicates so that differential effects of the two loci were not averaged out in the final analysis. Statistics were averaged over 50 replicate simulations for each particular set of parameter values.

## Results

### Trait divergence and genetic modularity under constraints to genetic decoupling

In the absence of genetic architecture evolution (*µ*_*pleio*_ = *µ*_*mutcor*_ = 0), traits are still capable of divergence, but do not follow trait optima closely since traits 3 and 4 get pulled away from their optima as traits 1 and 2 increase to follow theirs (Figure 5). With the introduction of variation in genetic architecture through mutation (*µ*_*pleio*_, *µ*_*mutcor*_ > 0), average trait values follow their optima more closely and the capability of trait divergence increases as mutation rates in genetic architecture increases, which leads to higher average population fitness values by generation 5000 (Figure 6). Also by generation 5000, simulations with higher pleiotropic connection mutation rates (*µ*_*pleio*_ *≥* 0.001 or *µ*_*mutcor*_ = 0.01) have distinctly modular genetic cor-relation structures with stronger correlations between traits 1 and 2 than between traits 3 and 4 (Figure 7). But at the highest pleiotropic connectivity mutation rate (*µ*_*pleio*_ = 0.01) the genetic integration of trait 3 with 4 (and even trait 1 with 2) is no longer as strong (i.e., the genetic correlation drops, see Figure 7). An increase in pleiotropic connectivity variation has a greater impact on trait divergence evolution and modularization of the genetic architecture of the traits than the same increase in mutational correlation variation, which is evident when either *µ*_*pleio*_ or *µ*_*mutcor*_ is the same as the allelic mutation rate (*µ* = 0.001). Even when the mutation rate for mutational correlation is at the highest tested (*µ*_*mutcor*_ = 0.01), an increase in the mutation rate for pleiotropic connections still improves the ability for traits to diverge, which can be seen in the decrease in variance over replicate simulations (Figure 5).

**Figure 5:**
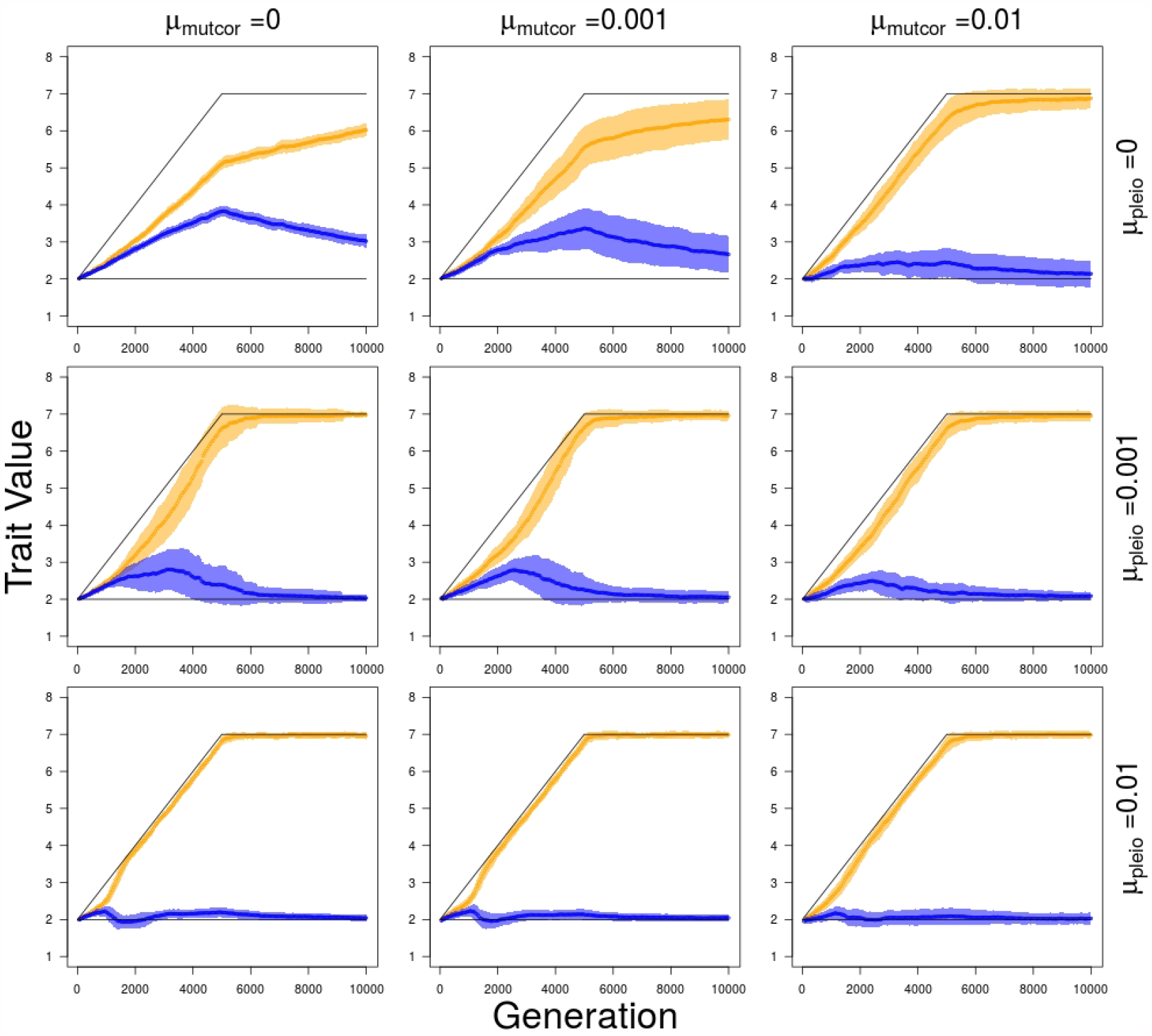
Trait value divergence over 5,000 generations of divergent selection on Traits 1 and 2 (Trait Module 1) followed by 5,000 generations of stabilizing selection for different combinations of mutation rate in pleiotropic connectivity (*µ*_*pleio*_) and mutational correlations (*µ*_*mutcor*_). Orange – average values of Traits 1 and 2; Blue – average values of Traits 3 and 4; Black – trait value optima for Trait Modules 1 and 2. Shaded regions show standard errors of the mean for 50 replicate simulations.

**Figure 6:**
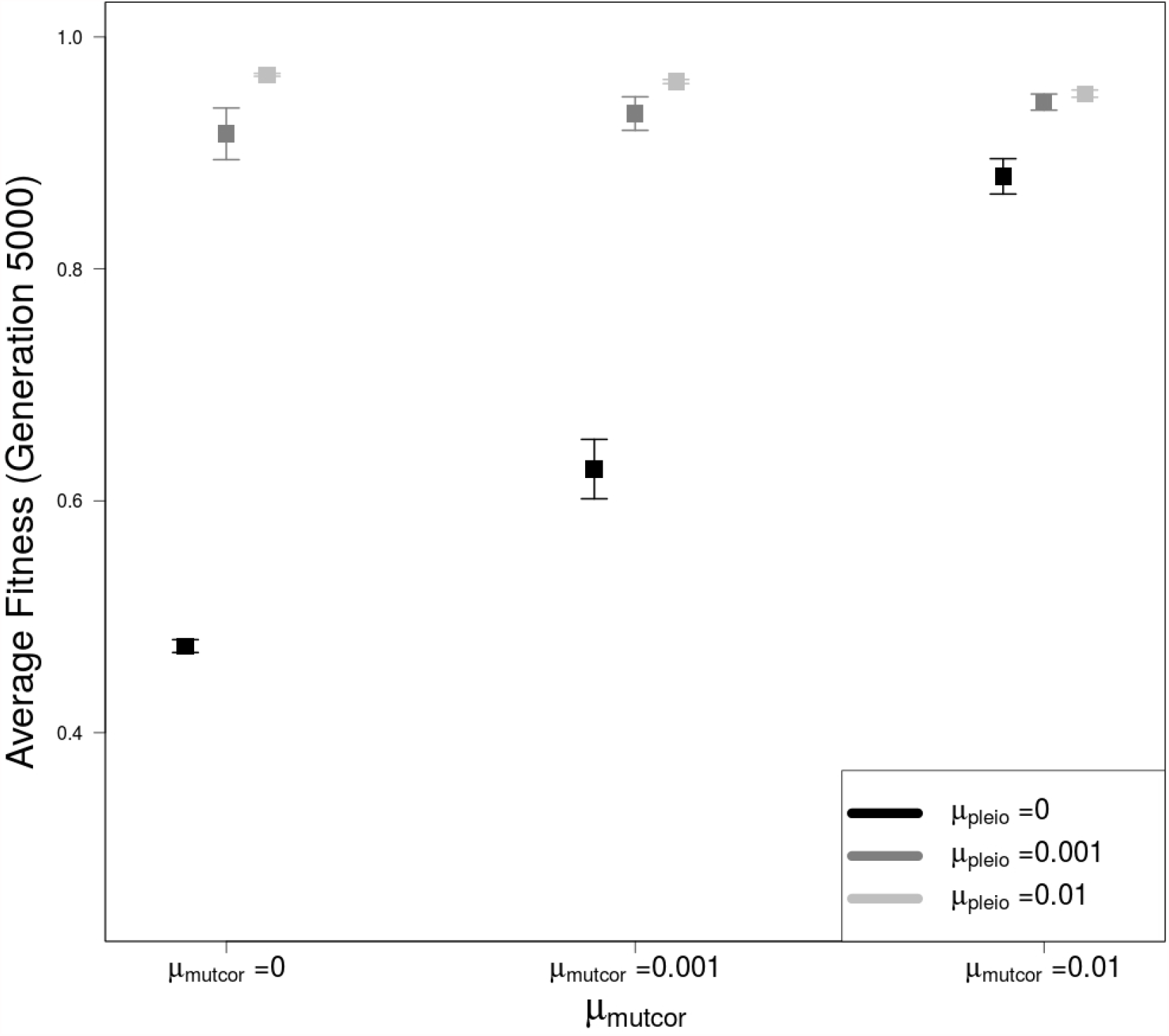
Average population fitness after 5,000 generations of divergent selection on Traits 1 and 2 (Trait Module 1) for different combinations of mutation rate in pleiotropic connectivity (*µ*_*pleio*_) and mutational correlations (*µ*_*mutcor*_). All error bars represent standard errors of the mean for 50 replicate simulations.

**Figure 7:**
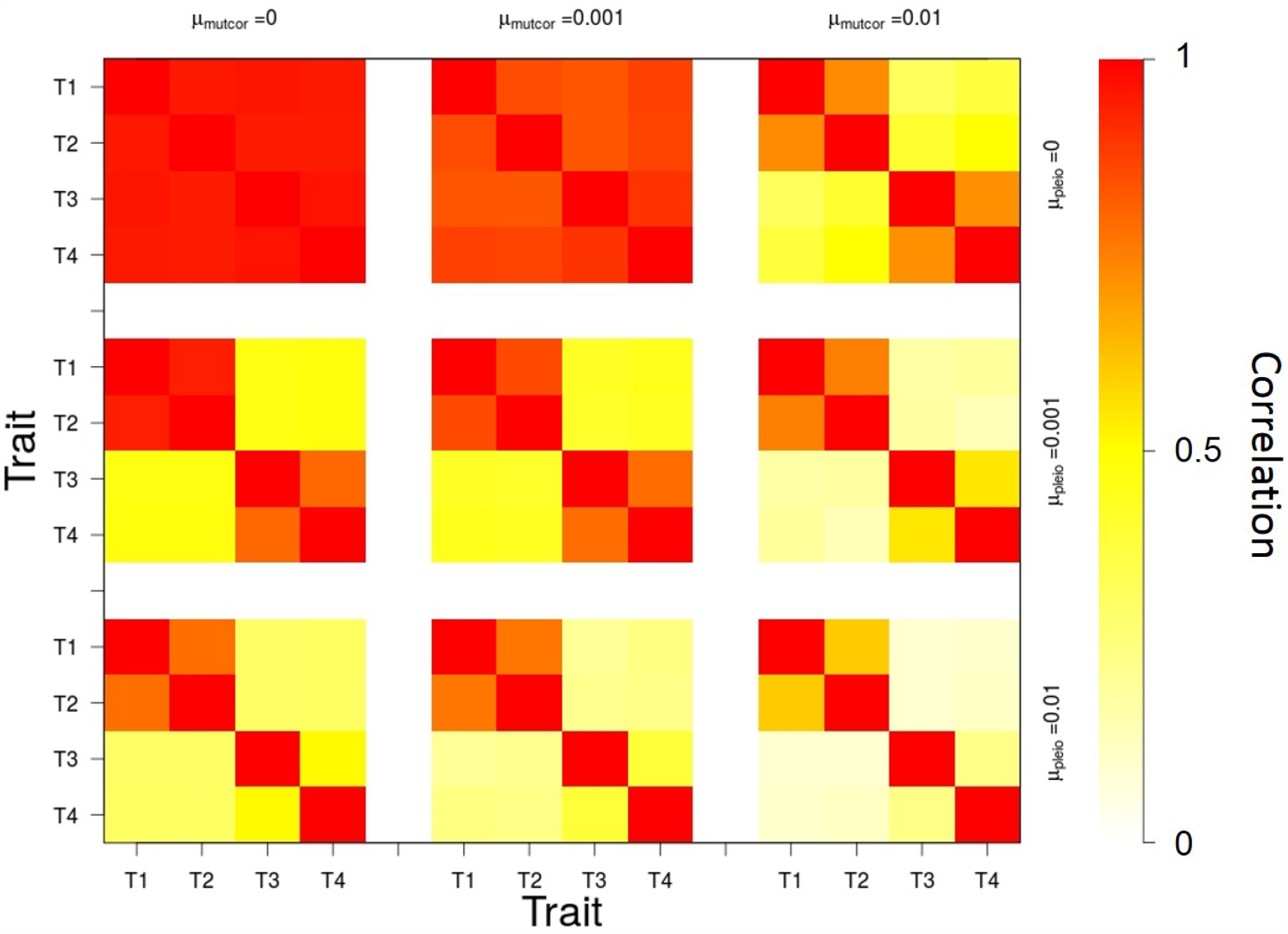
Genetic correlations between traits after 5,000 generations of divergent selection on Traits 1 and 2 (Trait Module 1) for different combinations of mutation rate in pleiotropic connectivity (*µ*_*pleio*_) and mutational correlations (*µ*_*mutcor*_). Red – higher genetic correlation. White – no genetic correlation

### Effects of pleiotropic connectivity and mutational correlation evolution on rate and extent of trait decoupling

When evolution of pleiotropic connections is possible (*µ*_*pleio*_ *>* 0), the most common allele in almost all cases is one that maintains connections to Traits 1 and 2, but has lost connections to traits 3 and 4 after two mutational events. This allele is found in Locus 1 or 2 at a frequency of 0.873 averaged over the populations of all simulations where evolution of pleiotropic connections is possible. The allele goes to fixation or near fixation in one locus where its pleiotropic degree decreases from four to two, and this happens more rapidly as *µ*_*pleio*_ increases (Figure 8). The decrease in pleiotropic degree resulting from the increase in frequency of this allele coincides with the modularization of genetic correlations, the divergence of traits and the increase in fitness. The proportion of times in which this particular allele becomes common in Locus 1 or in Locus 2 is approximately equal over all simulations (0.491 and 0.509, respectively, over 300 simulations) and is never observed in both loci in any one individual.

**Figure 8:**
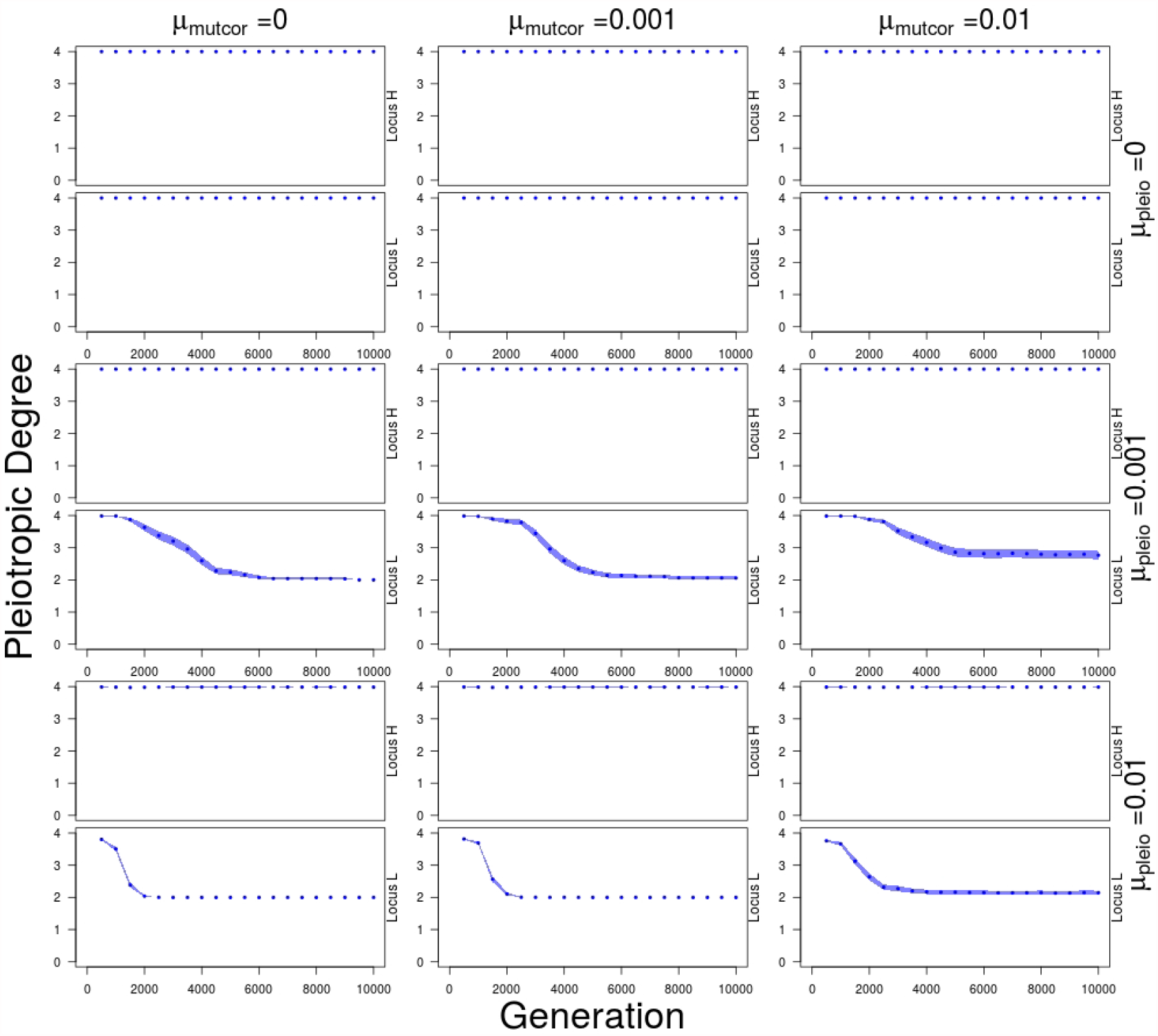
Average number of traits connected to each locus over 5,000 generations of divergent selection on Traits 1 and 2 (Trait Module 1) followed by 5,000 generations of stabilizing selection for different combinations of mutation rate in pleiotropic connectivity (*µ*_*pleio*_) and mutational correlations (*µ*_*mutcor*_). Loci are sorted so that locus with higher pleiotropic degree (Locus H) is always shown above and lower pleiotropic degree (Locus L) shown below. Shaded regions show standard errors of the mean for 50 replicate simulations.

When the mutation rate for pleiotropic connectivity (*µ*_*pleio*_) is zero, mutational correlation evolution can still lead to trait divergence but this takes longer, does not diverge as fully, and therefore leads to lower population mean fitness. Evolution of the mutational correlation occurs by a general decrease in all mutational correlations between traits at a rate determined by the mutation rate of mutational correlations (Figure 9). When the mutation rate at the mutational correlation loci is higher than the pleiotropic mutation rate then genotypic patterns do emerge where one locus disconnected from Trait 3 combines with lower mutational correlations between Traits 1 and 4 or 2 and 4, *or* a locus disconnected to Trait 4 combines with lower mutational correlations between Traits 1 and 3 or 2 and 3 (at frequencies of 0.16 and 0.10 over 50 replicates, respectively). But even in the case with a higher mutation rate for mutational correlation than the pleiotropic connectivity mutation rates, full subfunctionalization (one locus loses connections to Traits 3 and 4) is a possible outcome occurring in 18% of 50 replicates after 5,000 generations.

**Figure 9:**
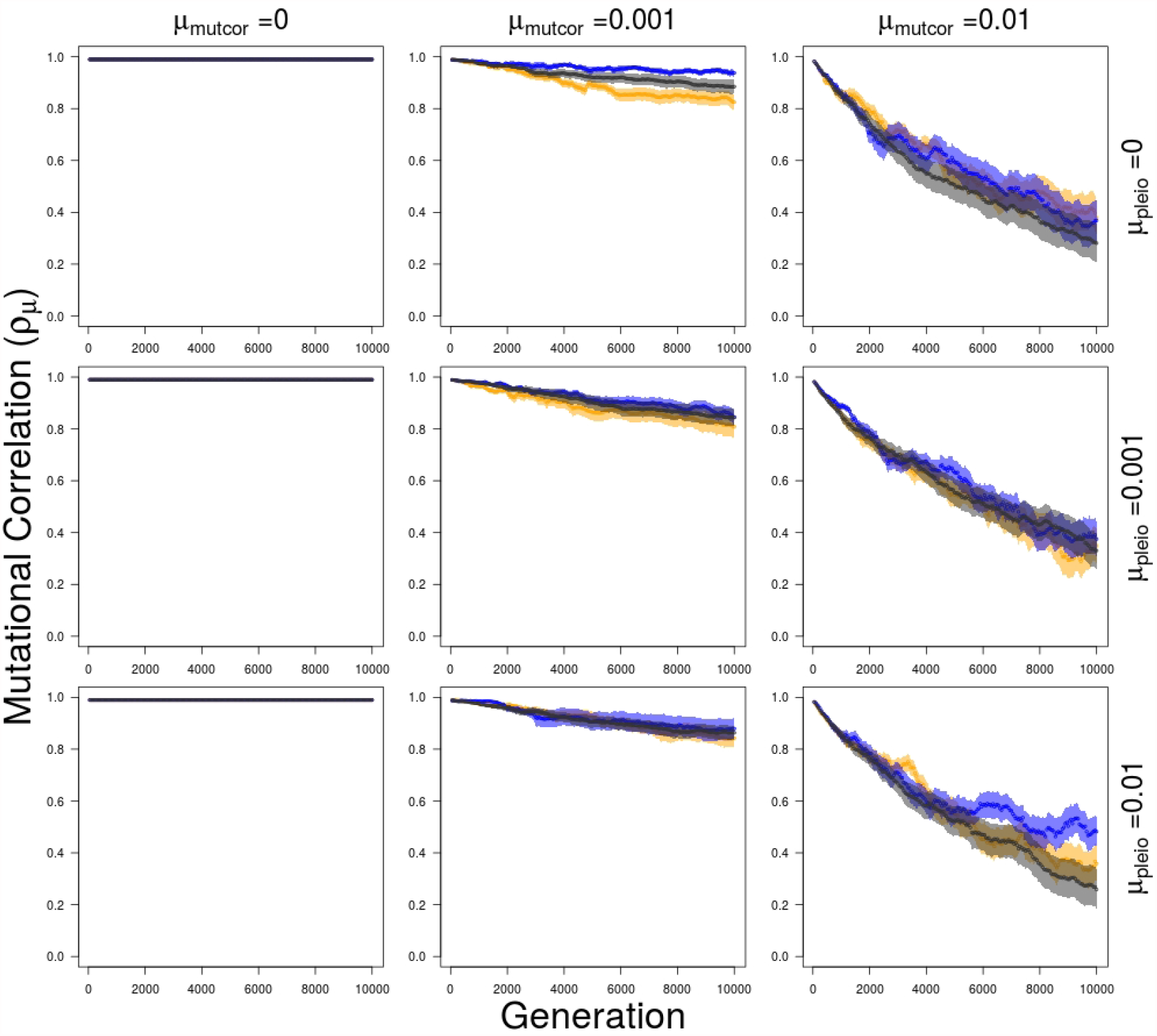
Average within and between trait module mutational correlation over 5,000 generations of divergent selection on Traits 1 and 2 (Trait Module 1) followed by 5,000 generations of stabilizing selection for different combinations of mutation rate in pleiotropic connectivity (*µ*_*pleio*_) and mutational correlations (*µ*_*mutcor*_). Orange – mutational correlation between Traits 1 and 2 (within Trait Module 1); Blue – mutational correlation between traits 3 and 4 (within Trait Module 2); Black – average mutational correlations between Traits 1 and 3, 1 and 4, 2 and 3, and 2 and 4 (between Trait Module 1 and 2). Shaded regions show standard errors of the mean for 50 replicate simulations.

### Effect of mutational correlation initial conditions set to zero (ρ_µ_ = 0 versus ρ_µ_ = 0.99)

In simulations where all mutational correlations are initialized at zero, there is little to no constraint on trait divergence despite full pleiotropic connectivity. This can be observed in trait values that follow their optima closely, leading to little reduction in fitness as optima for Traits 1 and 2 diverge from Traits 3 and 4, with little evolution in mutational correlations and pleiotropic degree during divergent selection (Figure 10). There are still patterns of genetic architecture evolution as alleles with lowered pleiotropic degree still emerge in the populations, but fixation is not common nor are any allelic patterns of mutational correlations.

**Figure 10:**
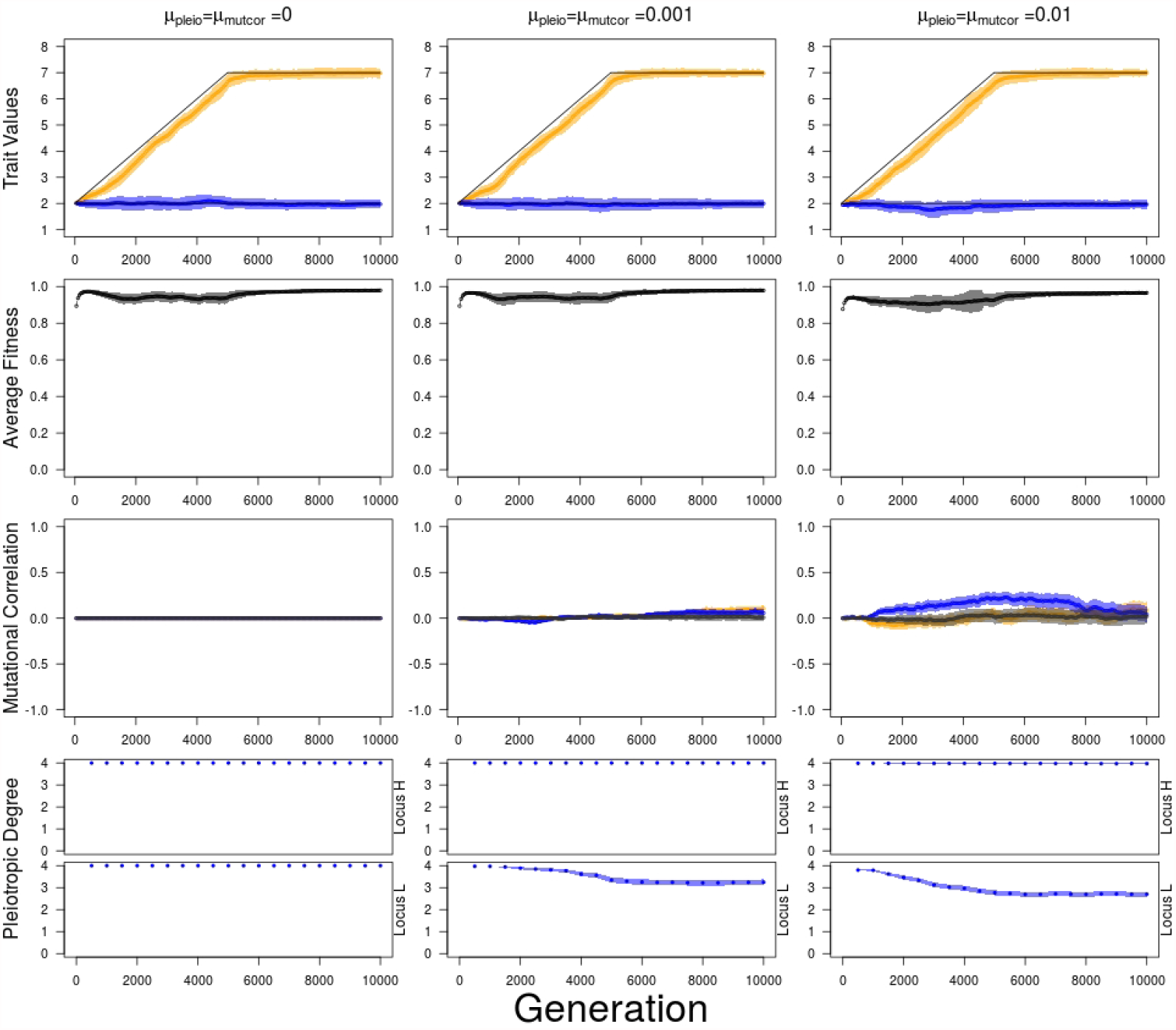
Average trait value, fitness, mutational correlation and pleiotropic degree, when all *ρ*_*µ*_ values are initialized to 0, over 5,000 generations of divergent selection on Traits 1 and 2 (Trait Module 1) followed by 5,000 generations of stabilizing selection for different combinations of mutation rate in pleiotropic connectivity (*µ*_*pleio*_) and mutational correlations (*µ*_*mutcor*_). For pleiotropic degree, loci are sorted so that locus with higher pleiotropic degree (Locus H) is always shown above and lower pleiotropic degree (Locus L) shown below. Orange – Trait 1 and 2 values or mutational correlation between Traits 1 and 2 (Trait Module 1); Blue – Trait 3 and 4 values or mutational correlation between Traits 3 and 4 (Trait Module 2); Black – average mutational correlations between Traits 1 and 3, 1 and 4, 2 and 3, and 2 and 4 (between Trait Modules 1 and 2). Shaded regions show standard errors of the mean for 50 replicate simulations.

## Discussion

### Evolution in pleiotropic connectivity and mutational correlation can lead to trait divergence

Previous models of genetic architecture evolution have shown that evolution in pleiotropic connections and mutational correlation can influence genetic correlation between traits and therefore responses to selection, but as far as we are aware this is the first time both have been allowed to evolve in the same model. When a genetic architecture is highly constraining to the decoupling of some traits from others, then evolution of the structure of the genetic architecture itself can clearly facilitate the rate and extent of trait divergence. Although genetic architecture may evolve through changes in pleiotropic connectivity between genes and traits, and in the mutational correlations between traits, the former leads to a greater release of genetic constraints and faster adaptation in the corridor selection regime. A qualitative distinction exists between these two types of genetic constraints to decoupling for two reasons. First, genetic constraints based on mutational correlation distributions are more difficult targets of selection compared to pleiotropic connections because mutations on modifiers of genetic correlations do not affect the trait phenotypes directly, whereas a single allele that differs in its pleiotropic connectivity does. Second, mutational correlations require pleiotropic connections to be effectual on traits (there can be no mutational correlations if a QTL affects only one trait), whereas the latter can affect the rate of adaptation regardless of mutational correlation (Baatz and Wagner, 1997; Chebib and Guillaume, 2017).

The results of this study corroborate results from previous models of pleiotropic evolution. We observe that divergent selection in the form of the corridor model leads to modular genetic architecture with greater genetic correlations between traits *within* trait modules and lower correlations *between* trait modules. This was also the case in both Melo and Marroig (2015) and Pavlicev et al. (2011) under the corridor model. Unfortunately, it is unclear whether patterns of partial modular pleiotropy that were responsible for the emergence of genetic modularity in our study were also observed in these studies because they did not report the most common resulting genotypes after corridor selection. Melo and Marroig (2015) did however vary the mutation rate in pleiotropic connectivity (while keeping allelic mutation rate the same) and found that when *µ*_*pleio*_ was 10 times greater than *µ*, there were higher within and between trait module correlation compared to when *µ*_*pleio*_ and *µ* were the same. Though our results corroborate this relationship as well, we cannot deduce the state of the pleiotropic connections that led to those results in their simulations. Their study also did not include evolution in mutational correlations so it is not possible to do a comparison on the relative effects of mutational correlation and pleiotropic connectivity evolution on patterns of genetic modularity. Pavlicev et al. (2011) had a deterministic model with rQTL (modifier loci) that affected the correlations between traits directly instead of affecting the pleiotropic connections, making it difficult to compare patterns of partial modular pleiotropic connectivity. Jones et al. (2007) found “extreme” variation among replicates in the average mutational correlation observed when *ρ*_*µ*_ was capable of evolving, similar to what was observed in our study (as well as when simulations were run with the same parameter values as the Jones et al. (2007) study; Supplemental Figure S1). This variation of the evolution of mutational correlation is likely due to an unstable equilibrium in the adaptive landscape in which highly positive or negative mutational correlations have a selective advantage over mutational correlations closer to zero (Lande (1980); Zhang and Hill (2002); Jones et al. (2007); Supplemental Figure S2).

### Patterns of pleiotropy

What explains the emergence of one dominant genotype that was observed with one locus losing its connections to Traits 3 and 4, and the other locus maintaining full pleiotropy? When mutational correlations are strong, genetic modularization should arise so that mutational effects can increase Traits 1 and 2 values without also increasing Traits 3 and 4, (especially when stabilizing selection is strong compared to directional selection). If stabilizing selection had been weaker and/or directional selection been much stronger, then more loci affecting the traits would have increased the proportion of advantageous mutations allowing for divergence (Hansen, 2003). For the same reason, we don’t observe complete genetic modularization with one locus only connected to Traits 1 and 2, and the other only connected to Traits 3 and 4. With both loci contributing to Traits 1 and 2, there is more mutational input to increase their values, giving support to the idea that intermediate levels of genetic integration will maximize evolvability when pleiotropic effects are all positive (Hansen, 2003). Also, since we were interested in the evolution of genetic architectures allowing trait decoupling, we started our simulations with a highly genetically integrated, monomorphic population. This makes evolution in our model dependent on de-novo mutations and as traits diverged, the negative effects of pleiotropy on traits under stabilizing selection increased, leading to modularization in the genetic architecture. But, if we had simulated genetic architectures where the allelic mutation rate (*µ*) was high enough and/or selection acted on many loci with small effects, pleiotropy may not have been as constraining, and integrated genetic architectures (loci affecting all traits) could be more evolvable. Whether integrated or modular genetic architectures will evolve in response to divergent selection is dependent on the relative effects of mutation and selection on the different traits (Pavlicev and Hansen, 2011). This also would have been true if standing genetic variation had already existed in pleiotropic connectivity and mutational correlations in a population prior to divergent selection. We could imagine that many possible combinations of pleiotropic connectivity and mutational correlation alleles that allow for increased variation and reduced covariation between traits could also exist. In those scenarios, genetic modularization may not be associated with trait divergence.

The results we obtain in this study are also related to work done on the evolutionary fate of duplicated, pleiotropic genes (Ohno, 1970; Hahn, 2009; Innan and Kondrashov, 2010; Guillaume and Otto, 2012). Previous models describe the conditions under which both genes remained fully pleiotropic, which is expected to be favorable when there is selection for increased dosage as we had for traits 1 and 2 (Ohno, 1970). There is some empirical evidence of this in ribosomal RNA, histone genes, as well as amylase genes in humans with high starch diets (Zhang, 2003; Perry et al., 2007; Qian et al., 2010). Other models describe when one or both genes lose their connection to some traits, known as subfunctionalization, if there is a relaxation of selection after duplication (Force et al., 1999; Lynch and Force, 2000). Empirical evidence for subfunctionalization exists for vertebrate limb evolution, as discussed in the introduction, as well as pathway specialization in plants (Bomblies and Doebley, 2006; Des Marais and Rausher, 2008). Compared to models with selection for increased dosage, our model has selection only for higher values in Traits 1 and 2, whereas selection for increased values in all four traits is expected to maintain all pleiotropic connections. The difference compared to neutral models where subfunctionalization is the result is that in our model there is no relaxation of selection due to duplication and redundancy. In that case, Guillaume and Otto (2012) showed that the maintenance of pleiotropy in one gene and subfunctionalization in the other (the most common outcome in our simulations) is predicted when there is asymmetry in either the trait contributions to fitness or in the expression levels of the genes. The gene with higher expression was predicted to remain fully pleiotropic, with loss of pleiotropy in the second, less expressed gene. Our results fit very well with that later outcome, although the conditions were different. In Guillaume and Otto (2012), a fitness trade-off emerged from the competitive allocation of the gene product (amount of protein produced) between two traits under positive selection (i.e., increased allocation to one trait reduced allocation to the other trait). The fitness trade-off in our model arose from the corridor model of selection whereby increased additive contributions to Traits 1 and 2 via fully pleiotropic mutations with correlated allelic values trade-off negatively with Traits 3 and 4 under stabilizing selection. The trade-off is quickly attenuated when the mutational correlations between traits under divergent selection decreases. Mutation in pleiotropic connections of the QTL was nevertheless more efficient in breaking the constraint to trait divergence. It is also a more plausible mechanism since mutations changing a transcription factors’s access to transcription binding sites may cause a drastic change associated with a change in pleiotropic connectivity.

### Empirical evidence for mutational correlation and pleiotropy

The pleiotropic connections and mutational correlations in our model abstract out the types of molecular level changes that may lead to changes in genetic correlations between traits. Some examples of variation in pleiotropic connectivity come from empirical studies on transcriptional regulation. For example, expression of the *Tbx4* gene (described earlier) is required not only for hindlimb development but is also expressed in genital development (Chapman et al., 1996). Although the upstream enhancer of *Tbx4*, hindlimb enhancer B (or HLEB), is functional in both hindlimb and genital development in both mice and lizards, HLEB appears to have lost its hindlimb enhancer function in snakes due to mutations in one of the enhancer’s binding regions (Infante et al., 2015). A more recent example comes from two species of Drosophila the diverged only 500,000 years ago. *D. yakuba* has both hypandrial and sex comb bristles whereas *D. santomea* has only sex comb bristles (Rice and Rebeiz, 2019). Quantitative trait mapping crosses between the species and with *D. melanogaster* revealed that a single nucleotide change in a regulatory enhancer of the *scute* gene, which promotes bristle development, was responsible for *D. santomea* losing its hypandrial bristles and increasing its sex comb bristle number (Nagy et al., 2018). These examples provide evidence that mutations in DNA binding sites can affect a gene’s pleiotropic degree, allowing for evolution of trait decoupling.

Correlated mutational effects, on the other hand, may arise from mutations that cause correlated effects in more than one of a gene’s molecular functions or from mutations causing correlated effects in a gene product’s multiple processes, but empirical data is still needed to discover the mechanisms underlying mutational correlations (Hodgkin, 1998; Wagner and Zhang, 2011). Even if the specific molecular mechanism that is the cause of correlation is not known, it is still possible to estimate the genomic **M**-matrix which describes the combined pattern of (co)variation arising from mutations in all loci that affect the traits of interest. Mutation accumulation experiments in *D. melanogaster* (Houle and Fierst, 2013) or *C. elegans* (Estes et al., 2005) provide examples of such genomic **M**-matrix estimates and show the existence of strong mutational correlation among morphological and life-history traits. Additionally, mutational correlations in *C. elegans* seem to correspond to phenotypic correlations among traits after removing environmental correlations and suggest that pleiotropy is somewhat restricted within traits of related function (Estes et al., 2005). Unfortunately, the **M**-matrix is only a summary statistic, which represents patterns of mutational variance across traits. It does not necessarily represent the correlations of mutational effects underlying that mutational variance between traits, which may be hidden due to multiple effects cancelling each other out.

It is also possible to discover evidence of modular pleiotropy from genome-wide studies using gene knock-out/-down experiments as was performed in yeast (Dudley et al., 2005; Güldener et al., 2005; Ohya et al., 2005), *C. elegans* (Sönnichsen et al., 2005), and the house mouse (Bult et al., 2008), which have shown that whole-gene pleiotropy is variable (not all genes affect all traits) and often modular (Wang et al., 2010; Wagner and Zhang, 2011). QTL studies further show variable pleiotropy in *D. melanogaster* (Mezey et al., 2005), threespine stickleback (Albert et al., 2008), the house mouse (Cheverud et al., 1997; Kenney-Hunt et al., 2008; Miller et al., 2014), and *A. thaliana* (Juenger et al., 2005), among others (Porto et al., 2016).

One empirical study based on human patient data manages to link mutational correlation with modular variation of pleiotropy by measuring both the genomic **M**-matrices and the pleiotropic degree of main and epistatic effects of mutations affecting the replicative capacity (fitness) of HIV-1 in different drug environments (Polster et al., 2016). In doing so, they discovered that epistasis can affect the pleiotropic degree of single mutations

producing modular genetic architectures and that epistatic-pleiotropic effect modules matched modules of fitness co-variation among drugs. These results suggest that epistasis may be fundamental in shaping the genetic integration itself, which may allow organisms to enhance their evolvability in the face of selection (Pavlicev et al., 2008, 2011; Pavlicev and Cheverud, 2015).

## Conclusion

Both pleiotropic connectivity and mutational correlation can constrain the divergence of traits under divergent selection, but when both can evolve, trait divergence occurs because pleiotropic connections are broken between loci and traits under stabilizing selection. The evolution of pleiotropic connectivity is favoured because it is an easier target of selection than a distribution of mutational effects. The most commonly observed genotype thus includes one locus that maintains connections to both traits under directional selection and both traits under stabilizing selection, and the other locus losing its connection to the traits under stabilizing selection (subfunctionalization). The subfunctionalization of one locus allows it to contribute to increasingly divergent trait values in the traits under directional selection without changing the trait values of the other traits, which leads to separate genetic modules. These results indicate that partial subfunctionalization is sufficient to allow genetic decoupling and the divergence of traits with little to no loss of average fitness.

## Acknowledgments

This manuscript benefited from the constructive comments provided by Thomas F. Hansen, Mihaela Pavlicev, Joachim Hermisson, Nicholas Barton, and an anonymous reviewer of the *Genetics* journal. J.C. and F.G. were supported by the Swiss National Science Foundation, grant PP00P3 144846 and PP00P3 176965 to F.G.

## Author Contributions

J.C. performed software modification for model implementation and acquisition of data, as well as drafting of manuscript. J.C. and F.G. performed study conception and design, analysis and interpretation of data, and critical revision of manuscript.

## Data Archival

The data and initialization files for this study are available online through Zenodo online repository at: https://zenodo.org/record/3980997#.XzPnFcBKi70 and code for simulations can be found at: https://github.com/jmchebib/nemo_evolving_pleio

## Conflict of interest disclosure

The authors of this article declare that they have no financial conflict of interest with the content of this article.

## Supplemental

**Figure S1:**
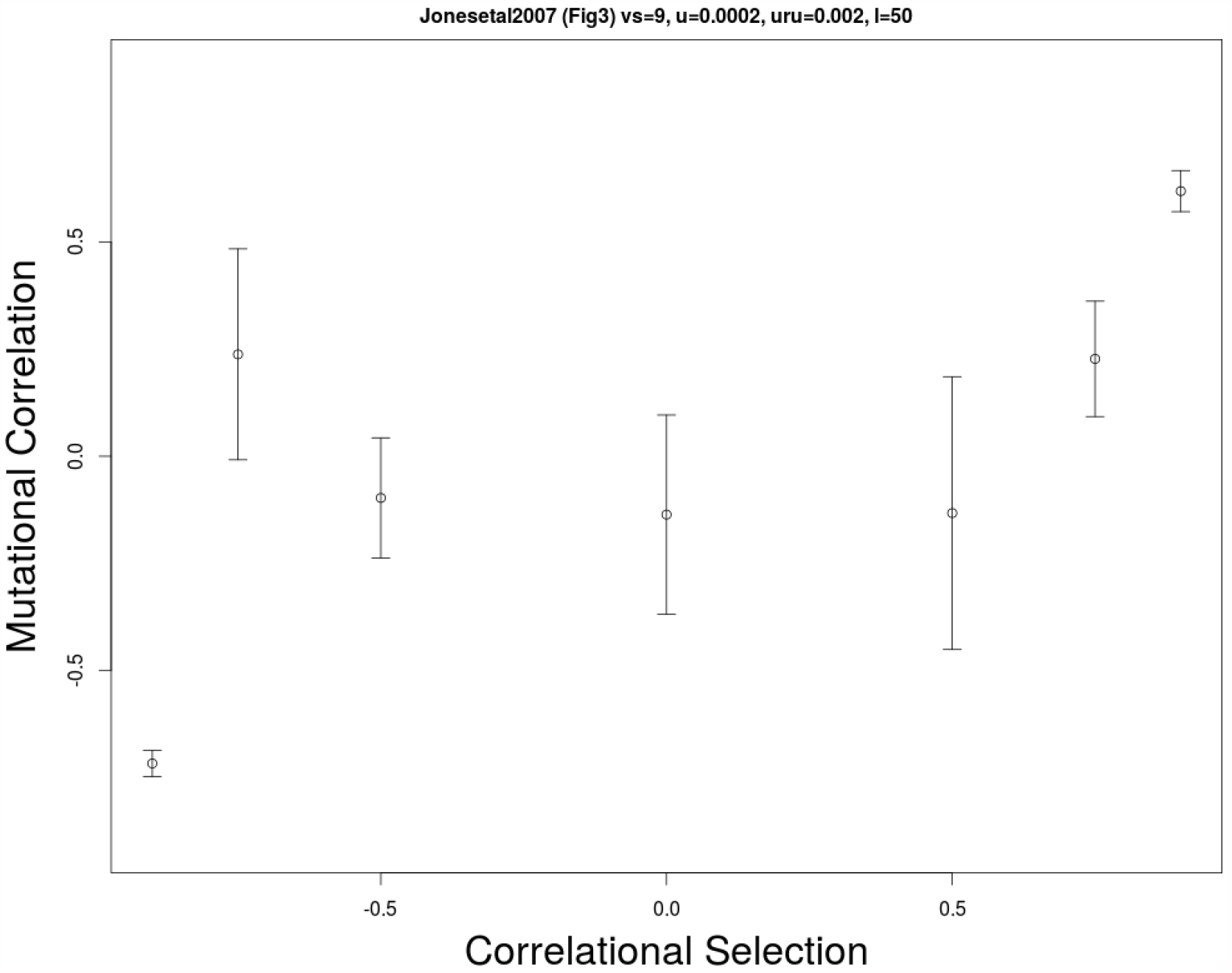
Average mutational correlation *ρ*_*µ*_ values for different values of correlational selection *ρ*_*ω*_ . Parameter values were chosen to match those used in Jones et al. 2007 wherever possible and averages were taken over values from every five generations after burn-in between generation 10,000 and 20,000. Number of loci = 50, Number of traits = 2, *N* = 2372, *ω*^2^ = 9, *µ* = 0.0002, *µ*_*mutcor*_ = 0.002, and *α*^2^ = 0.05.

**Figure S2:**
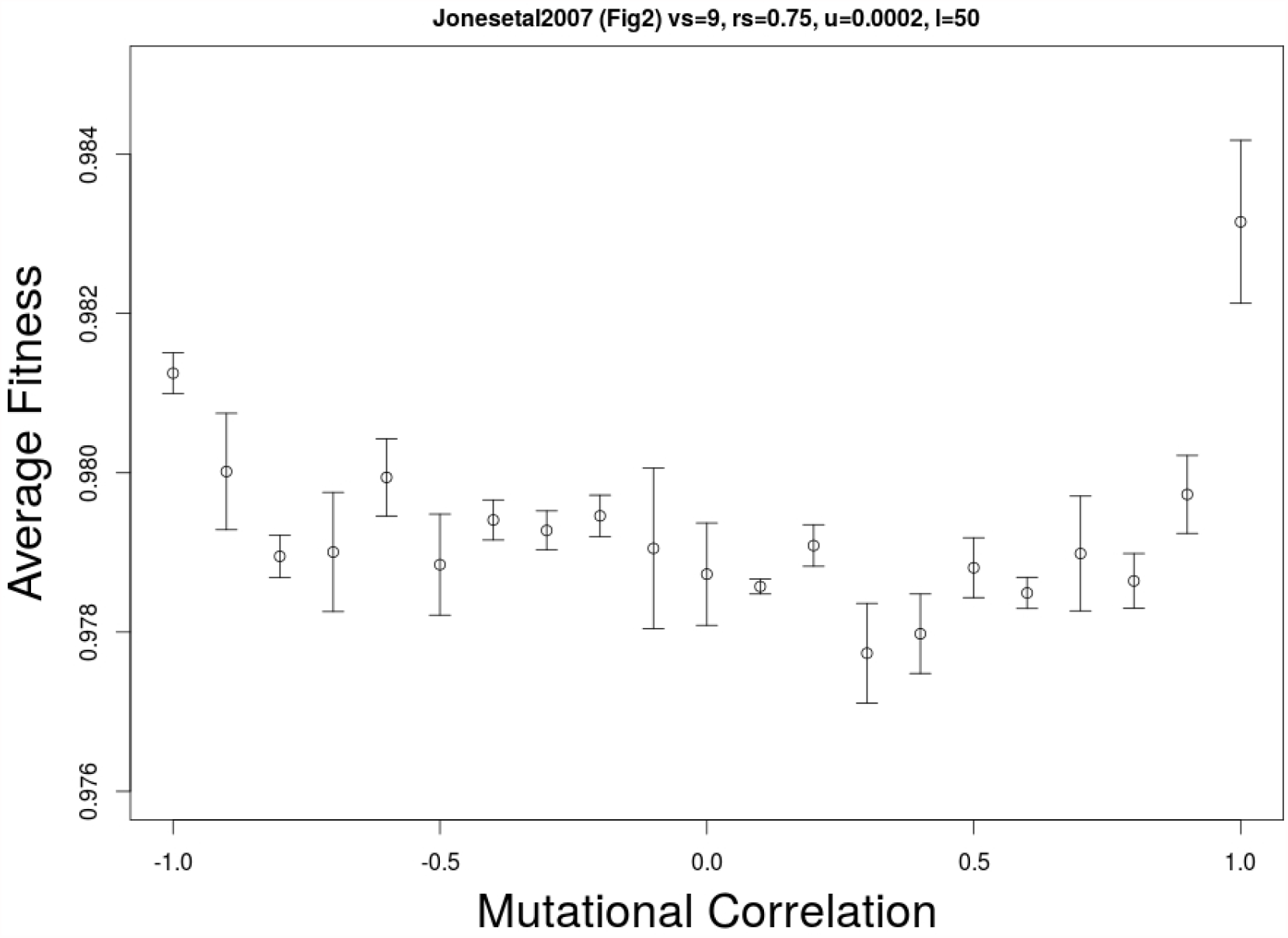
Average fitness for different values of mutational correlation (static). Parameter values were chosen to match those used in Jones et al. 2007 wherever possible and averages were taken over values from every five generations after burn-in between generation 10,000 and 15,000. Number of loci = 50, Number of traits = 2, *N* = 2372, *ω*^2^ = 9, *ρ*_*ω*_ = 0.75, *µ* = 0.0002, and *α*^2^ = 0.05.

## Notes

https://zenodo.org/record/3980997\#.XzPnFcBKi70

https://github.com/jmchebib/nemo_evolving_pleio

### Competing Interest Statement

The authors have declared no competing interest.

### Summary of Updates

Traits can diverge whether there is evolution in pleiotropic connectivity or mutational correlation but when both can evolve then evolution in pleiotropic connectivity is more likely to allow for divergence to occur. The most common genotype found is characterized by having one locus that maintains connectivity to all traits and another that loses connectivity to the traits under stabilizing selection (subfunctionalization). This is favoured because it allows the subfunctionalized locus to accumulate greater effect size alleles, contributing to increasingly divergent trait values in the traits under directional selection without changing the trait values of the other traits (genetic modularization).

